# SAMHD1 controls nucleoside-based cancer therapeutics, deoxyguanosine toxicity, and inflammation

**DOI:** 10.1101/2022.08.25.505278

**Authors:** Michael D. Moore, Errin Johnson, Chloe Jacklin, Steven Johnson, Owen Green, William James

**Affiliations:** Sir William Dunn School of Pathology, University of Oxford, South Parks Road, OX1 3RE; Department of Earth Sciences, University of Oxford, South Parks Road, OX1 3AN

**Keywords:** SAMHD1, dNTP, dG, toxicity, inflammation, uric acid

## Abstract

SAMHD1 is a key regulator of dNTP metabolism, being activated by dGTP and converting dNTPs into their respective nucleosides. As a result, it plays a significant role in several distinct diseases. Not only is SAMHD1 a restriction factor for HIV-1 replication, which requires dNTPs for replication, but it is also required for genome stability, is mutated in multiple cancers, and loss of function mutations cause Acardi-Goutières Syndrome, an early-onset inherited encephalopathy. In this paper we knock out the SAMHD1 gene in pluripotent stem cells, and show that it also affects nucleoside analogue sensitivity of dividing cells and deoxyguanosine toxicity in a cell cycle-independent manner. Comparing wild type and gene-edited macrophages, we also show that deoxyguanosine toxicity is associated with pro-inflammatory cell death, modifications in uric acid production and the formation of monosodium urate crystals within the macrophages. These novel features of the SAMHD1^−/−^ phenotype provide unique insights into the aetiology of Acardi-Goutières Syndrome, and have implications for certain chemotherapeutic agents.

## Introduction

The metabolism of dNTPs is a tightly regulated process, with multiple feedback mechanisms that provide homeostatic control(1). Sufficient levels of dNTPs are required for completion of DNA replication during the S phase of the cell cycle, but alterations in the balance between the different dNTPs can result in mutagenesis(2). In terminally differentiated cells, dNTPs are only required for DNA repair and mitochondrial DNA replication, and are kept to a minimum e.g. dNTPs in macrophages are 130-250-fold lower than in activated T cells(3). Non-dividing cells maintain low levels of dNTPs by regulating two pathways: by restricting de novo synthesis of dNTPs from rNTPs by ribonucleotide reductase (RNR) combined with catabolism of dNTPs by the nucleotide triphosphate triphosphohydrolase enzyme, SAMHD1(4). Thus, it is not surprising that mutations in key genes involved in dNTP metabolism, such as SAMHD1, are linked to mitochondrial DNA deletion disorders(5,6). Keeping dNTP concentrations low in non-dividing cells has the added benefit of restricting the replication of parasites (e.g. Leishmania and HIV-1) that require host dNTP to replicate their genome(3,7). Accordingly, SAMHD1 is a target for both pathogens and cancer, both of which benefit from loss of SAMHD1 function and the resultant elevation in dNTP levels. For example, SIV and HIV-2 encode VPx, which targets SAMHD1 for degradation, thereby allowing replication in non-dividing macrophages(8). Additionally, SAMHD1 mutations are frequent in numerous cancers, potentially as driver mutations(9,10). Importantly, SAMHD1 activity can impact the effectiveness of the nucleoside analogues with which both HIV and cancers are often treated(11,12). These are administered as prodrugs that require phosphorylation for activity, a process potentially countered by SAMHD1. The efficacy of different chemotherapeutic drugs within this class can vary widely depending on the type of cancer targeted. For example, cytarabine (Ara-C) shows very good response rates to haematological cancers such as lymphomas but is inactive against other “solid” tumours. Conversely Gemcitabine shows good activity against a variety of cancers.

Defects in metabolic control that result in imbalances in dNTP levels can have significant but often varied clinical outcomes, depending on the specific gene mutation. For example, severe combined immunodeficiency can result from elevated dATP due to mutations in adenosine deaminase (ADA), resulting in loss of both T and B lymphocytes(13). Immunodeficiency can also result from elevated dGTP due to mutations in purine nucleoside phosphorylase (PNP), resulting in loss of T cells and autoimmune complications such as lupus erythematosus(14). These clinical findings suggest that T cells are particularly sensitive to dNTP imbalance, a feature thought to be linked to their rapid cell division. Another disease linked with the salvage pathway of nucleotide metabolism, Lesch-Nyhan syndrome, is associated with neurological complications and severe gout due to loss of Hypoxanthine-guanine phosphoribosyltransferase (HGPRT), which results in excessive purine catabolism and elevated uric acid levels(15). Additionally, deficiency in SAMHD1 activity is one of the causes of an early onset encephalopathy, Acardi-Goutières Syndrome (AGS), which is associated with chilblain lupus, intracerebral calcifications, and mental retardation(16–18). In this study, we hypothesize that loss of function of SAMHD1 results in altered drug sensitivity and purine metabolism.

## Results

### Drug sensitivity of cell lines do not correlate with their SAMHD1 levels

Upon T cell activation, SAMHD1 activity is repressed, both by downregulation of expression and phosphorylation at T592 of the protein(19), in order for sufficient levels of dNTPs to be produced for cell proliferation(20,21). This feature of T cell activation is not only exploited by HIV-1 but can also be observed in T cell cancers and cell lines derived from them, which often express little to no SAMHD1 (e.g. the CEM T cell line shown in Figure 1A). This contrasts with non-dividing cells, such as those of the myeloid lineage, that express high levels of unphosphorylated SAMHD1, as seen in the pluripotent stem cell-derived macrophages (pMac, Figure1A). Proliferation of cells derived from the myeloid lineage, such as the THP1 cell line, which derives from a monocytic leukemia, is associated with phosphorylation of SAMHD1 (Figure 1A).

**Figure 1.**
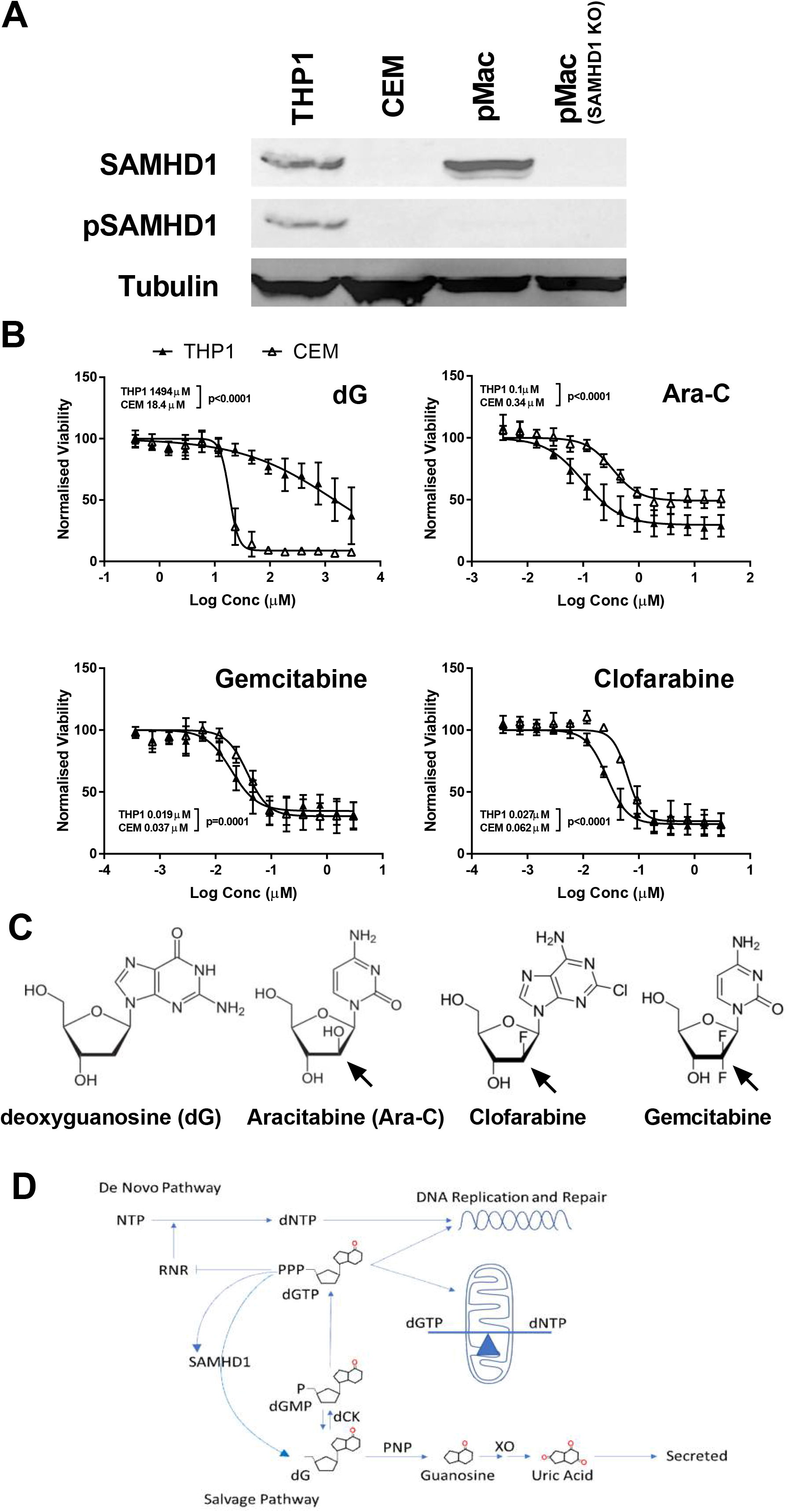
Cancer cell line response to chemotherapy is not consistent with SAMHD1 expression profiles. (A) Western blot showing the levels of total SAMHD1 and phosphorylated SAMHD1 in the monocytic THP1 cell line, T lymphoblast CEM cell line, pluripotent stem cell derived macrophages (pMac) and the SAMHD1 knockout macrophages (SAMHD1 KO) compared with the tubulin loading control. (B) Viability assays for THP1 (closed triangles) and CEM (open triangles) cells exposed to increasing concentrations of four drugs. Results are plotted as mean +/- standard deviation (SD) of relative viability to an untreated control. P-values represent the statistical significance for the EC50 measurements being different (F-test). (C) Molecular structures of the four drugs used in the study with an arrow highlighting the major alterations from a standard deoxyribose ring. (D) Schematic of dGTP metabolism under physiological conditions.

The impact of SAMHD1 on nucleoside-based cancer therapeutics was initially tested by comparing the sensitivity of CEM and THP1 cells to three different drugs that have been used clinically (Ara-C, Gemcitabine and Clofarabine) and a natural nucleoside with cytotoxic properties (deoxyguanosine or dG)(22) (Figure 1B+C). The two cell types demonstrated differing sensitivity to the drugs, particularly to dG, with the cells lacking SAMHD1 being 80-fold more sensitive to dG (Figure 1B). Deoxyguanosine enters the salvage pathway for nucleotide metabolism and is phosphorylated to dGTP for use in DNA replication and repair (Figure 1D). Elevated levels of dGTP are cytotoxic, through an incompletely understood pathway, though dGTP is known to allosterically inhibit de novo dNTP production and be degraded by SAMHD1. In contrast to dG, all other drugs showed greater toxicity towards THP1 cells. The lack of consistency suggests that either SAMHD1 is not involved in regulating drug sensitivity, or additional metabolic changes resulting from the extreme aneuploidy of cancerous cell lines makes meaningful comparisons impossible.

### Drug sensitivity of pluripotent stem cells is controlled by SAMHD1

In contrast to the cell lines, pluripotent stem cells have a normal karyotype. Thus, when genetic modifications are made using technology such as CRISPR-Cas9, the impact of the modification can be assessed within the background of normal cellular function. Pluripotent stem cells express very low levels of SAMHD1, with a high level of phosphorylation. Nevertheless, we hypothesised that loss of any residual activity of SAMHD1 would result in enhanced sensitivity to cytotoxic nucleoside analogues. Using a previously generated SAMHD1 knockout line (23) and its isogenic parental line we measured the cytotoxic profiles of the same panel of drugs (Figure 2A). The sensitivities of these cells provided a much more consistent set of data demonstrating the clear effect that SAMHD1 activity has on cytotoxicity of the drugs. Both dG and Ara-C were >15 fold more toxic in SAMHD1-knockout cells compared with wild-type cells. Clofarabine activity was enhanced 6-fold by loss of SAMHD1, whereas Gemcitabine was only marginally affected. The reproducibility of these results was confirmed for dG and Ara-C in three independent wild-type backgrounds and with three SAMHD1 knockout clones (Supplementary Figure 1A+B). The cytotoxic potency of the nucleoside drugs on SAMHD1 wild type versus knock-out stem cells corresponds to the molecular deviation of the drugs’ ribose ring from that of the natural substrates of SAMHD1, dNTPs. Deoxyguanosine is a natural nucleoside (Figure 1C), hence it is unsurprising that its toxicity is limited by SAMHD1, whose activity would reduce the levels of the active metabolite dGTP by conversion back to dG. Ara-C contains an arabinose ring rather than a deoxyribose ring, which resembles a ribose ring but with the 2’OH group in an altered stereochemical location (Figure 1C). This alteration sterically hinders the rotation of DNA when the triphosphate of Ara-C is incorporated into the growing strand and results in DNA replication failure. However, it would appear that SAMHD1 is capable of accommodating the additional altered 2’OH group, as it is able to limit the toxicity of Ara-C by preventing the accumulation of the toxic Ara-C triphosphate. Clofarabine and Gemcitabine also differ from natural nucleosides at the 2’ position, incorporating one or two fluorines, respectively. The SAMHD1 sensitivity profiles of these two drugs suggest that the active site of SAMHD1 is not accessible to fluoro-modified ribose, and in particular does not accommodate two fluorines. Using in silico molecular modelling, it would appear that while no significant clashes are seen within the active site of SAMHD1, there is the potential for significant electrostatic repulsion between fluorines at the 2’ position and the pi-electrons of Tyr374 in SAMHD1 (Supplementary Figure 1C+D), which may prevent the triphosphates of Clofarabine and Gemcitabine from being efficiently catabolised by SAMHD1(24,25).

**Figure 2.**
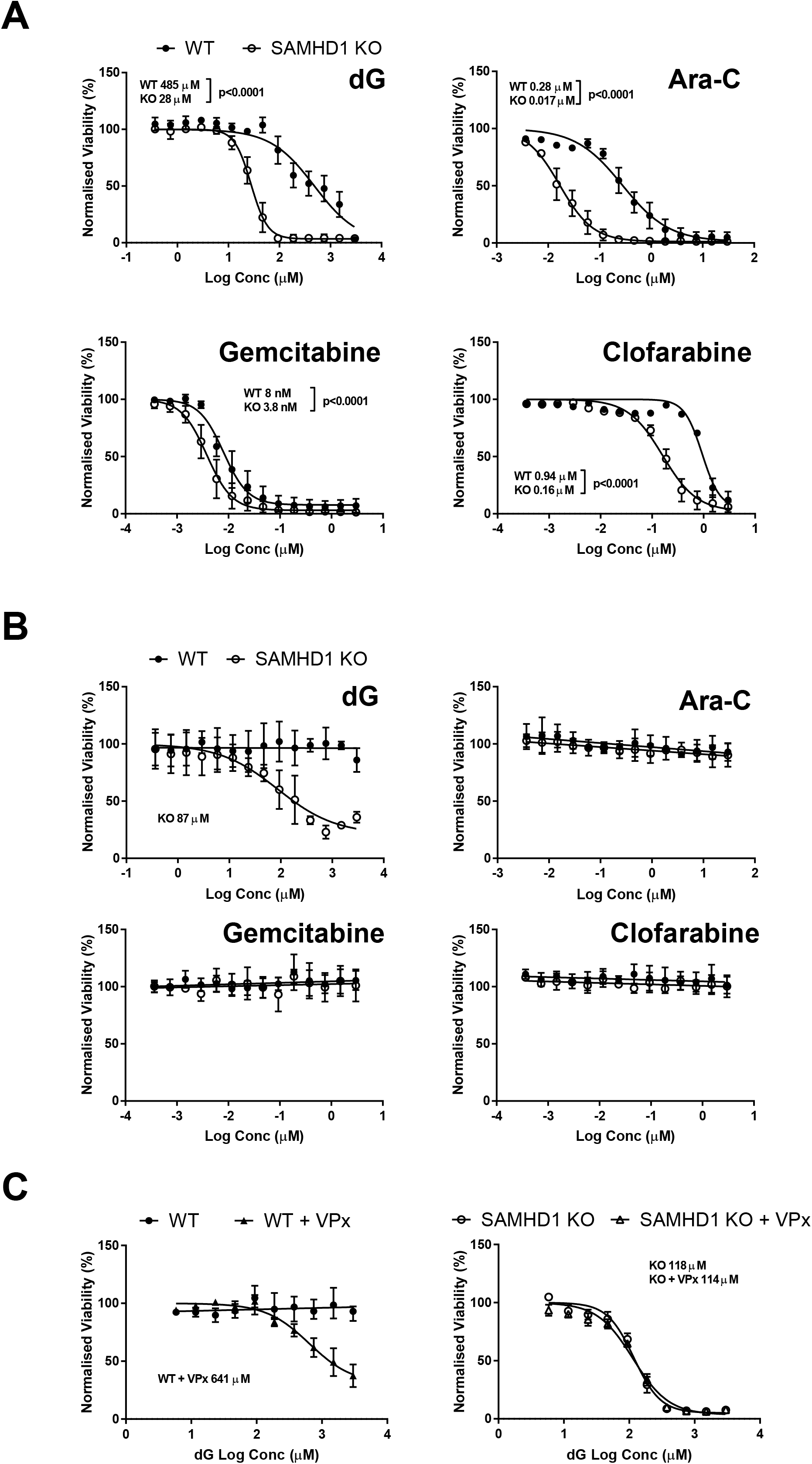
Drug dependent effects of SAMHD1 on cytotoxicity. (A) Viability assays for wild-type (WT) pluripotent stem cells (closed circles) against an isogenic knockout clone for SAMHD1 (open circles) in the presence of increasing concentrations of drugs. Results are plotted as mean +/- SD of the relative viability to an untreated control sample. P-values represent the statistical significance for the EC50 measurements being different (F-test). (B) As for (A) but using pluripotent stem cell-derived macrophages. (C) Viability assays of stem cell-derived macrophages in the presence of increasing quantities of dG, in the absence (circles) or presence (triangles) of VLPs carrying the SIV protein VPx, known to degrade SAMHD1.

### Nucleoside analogue activity is cell cycle dependent

Although uncontrolled cell division is a hallmark of cancer, non-dividing cells, such as quiescent cancer stem cells and tumour-associated macrophages (TAMs), are involved in cancer development(26,27). To observe the effect of SAMHD1 activity on the panel of drugs in authentic non-dividing cells we differentiated the isogenic pair of pluripotent stem cells into pMacs(28). We have previously demonstrated that pMacs are terminally differentiated, c-Myb-independent and, therefore, are an accurate physiological and ontological model of tissue-resident macrophages(29). Exposure of these non-dividing cells to the antiproliferative chemotherapeutic agents Ara-C, Gemcitabine and Clofarabine unsurprisingly had no impact on cell viability (Figure 2B). However, increasing levels of dG was toxic to pMacs, exclusively in the absence of SAMHD1 (EC50 in KO 87 μM). This suggests that an imbalance of dNTPs is toxic in a cell cycle-independent manner. These results were confirmed by treating both the control and knockout cells with virus-like particles (VLPs) containing the SIV protein, VPx, which is known to degrade SAMHD1 and enable SIV infection of macrophages(8). Upon exposure of the wild-type pMacs to VPx, they became sensitive to elevated levels of dG (EC50 640 μM), whereas, as expected, VPx had no effect on the SAMHD1 knockout cells (Figure 2C and Supplementary Figure 2A).

### Deoxyguanosine exposure alters purine metabolism

An interesting feature of exposure of wild-type macrophages (either stem cell-derived or blood monocyte-derived) to high levels of dG was the development of large, dark inclusion bodies by light microscopy, not observed in control cells (Figure 3A). Observation of dG-treated cultures by scanning electron microscopy compared with control cells identified disrupted cellular morphology, with localised membrane contortions and crystalline non-cellular debris (Figure 3B) that could be found within the remains of a dead cell (Supplementary Figure 3A, left) or being extruded from a living cell (Supplementary Figure 3A, right). Using transmission electron microscopy, the inclusion bodies could be seen as either enlarged circular vesicles or defined needle like structures, neither of which were detected in control cells (Figure 3C and Supplementary Figure 3B, left). Multiple forms of the large vesicles were observed, either with or without an intact membrane and with either fibrous or crystalline contents, which may represent various stages of crystal development (Supplementary Figure B, right).

**Figure 3.**
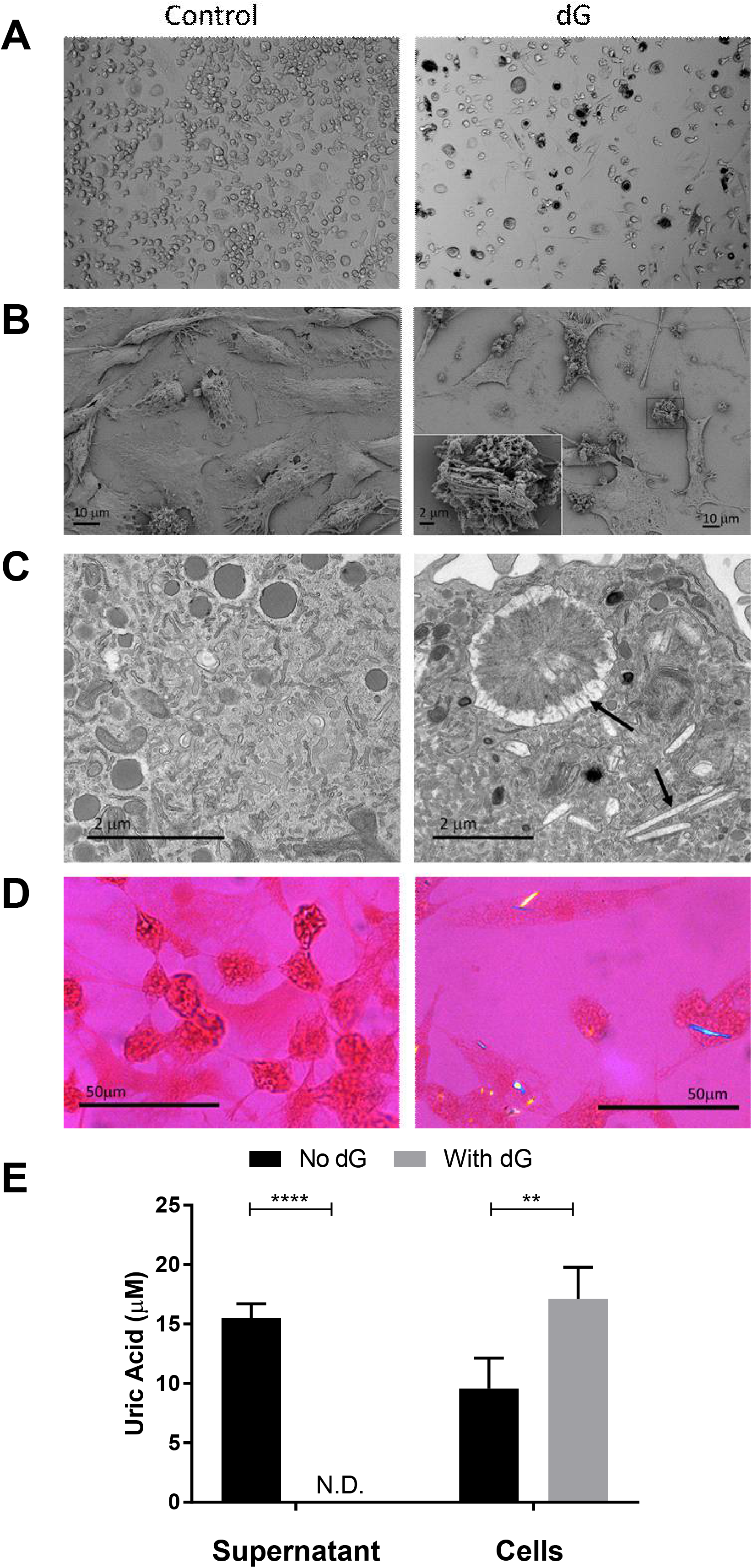
Addition of dG to macrophage cultures results in uric acid crystal formation. (A) Light microscopy of monocyte-derived macrophages in the absence (Control) or presence of 1mM dG (after 7 days’ exposure). (B) SEM images of untreated pMac or pMac treated with 1.5 mM dG for 8 days. (C) TEM images of (A) with needle-like crystals and enlarged vesicles highlighted with arrows. (D) pMacs with and without dG for 5 days observed under a polarised microscope with a gypsum plate to identify birefringent crystals, seen within the dG treated cells as orange and blue needles. (E) Uric acid levels within the supernatant of untreated (No dG) or 3 mM dG treated (with dG) pMacs 5 days post treatment. (****p<0.0001, **p=0.0029, 2-way ANOVA with Sidak’s multiple comparison test).

Deoxyguanosine is a purine nucleoside, and given that the end product of purine catabolism is uric acid in humans (Figure 1D), we hypothesised that the inclusion bodies were comprised of monosodium urate (MSU) crystals. These are needle-like crystals found within the synovial fluid of gout sufferers that show a characteristic birefringence under polarised light(30). Using a polarised light microscope and gypsum plate filter we were able to identify numerous birefringent crystals within the pMacs exposed to dG but not of control cells (Figure 3D, right versus left panel). To confirm the presence of elevated uric acid after treatment of pMacs with dG, we performed a uric acid assay on the cell supernatant and on the cells. Interestingly, pMacs under physiological conditions secrete a high level of uric acid into the supernatant, but upon treatment with dG this is prevented (Figure 3E). Conversely, higher levels of uric acid were recovered from the cells after exposure to dG (Figure 3E), a component of which is presumably contained within the cells as monosodium urate crystals (Supplementary Figure 5A).

### SAMHD1 activity alters the impact of deoxyguanosine on purine metabolism

Having shown that dG treatment is linked to uric acid production in wild-type cells, we hypothesised that loss of SAMHD1 activity would not only make the cells more sensitive to dG toxicity but would also alter the levels of uric acid secretion, both at steady state and after exposure to dG (Figure 1D). Therefore, we measured the levels of uric acid in the supernatant of wild-type and SAMHD1-knockout pMac cultures in the presence of increasing concentrations of dG. To simulate the loss of SAMHD1 in the wild-type control cells we also treated both control and knockout cells with VPx-containing VLPs. The results in Figure 4A confirm the impact of dG on secretion of uric acid into the supernatant of wild-type pMacs, previously noted in Figure 3E. A similar effect can be seen in SAMHD1 knockout cells (Figure 4B), with loss of uric acid secretion as dG increases, however this occurs at lower levels of dG, in line with the increased sensitivity of these cells to dG-mediated toxicity. In agreement with our hypothesis, we observed that SAMHD1-knockout pMacs secrete less uric acid into the supernatant than wild-type cells, a feature that is replicated by adding VPx to wild-type pMacs, but not SAMHD1-knockout pMacs, demonstrating it is a SAMHD1-dependent effect. Both the impact of dG and loss of SAMHD1 on uric acid production from pMacs was shown to be significant after performing multiple experiments at a single concentration of dG (Figure 4C+D) and was consistent across different donor macrophages (Supplementary Figure 2B). Why SAMHD1 loss results in lower levels of uric acid secretion can be explained by an alteration in the balance of the interconversion between dG and dGTP. Without SAMHD1 degrading dGTP to dG, under physiological conditions, dG would become limited. The downstream result of which would be reduced levels of guanosine produced from dG by PNP, and reduced uric acid from guanosine by Xanthine Oxidase (Supplementary Figure 5B), thereby reducing the amount of uric acid secreted.

**Figure 4.**
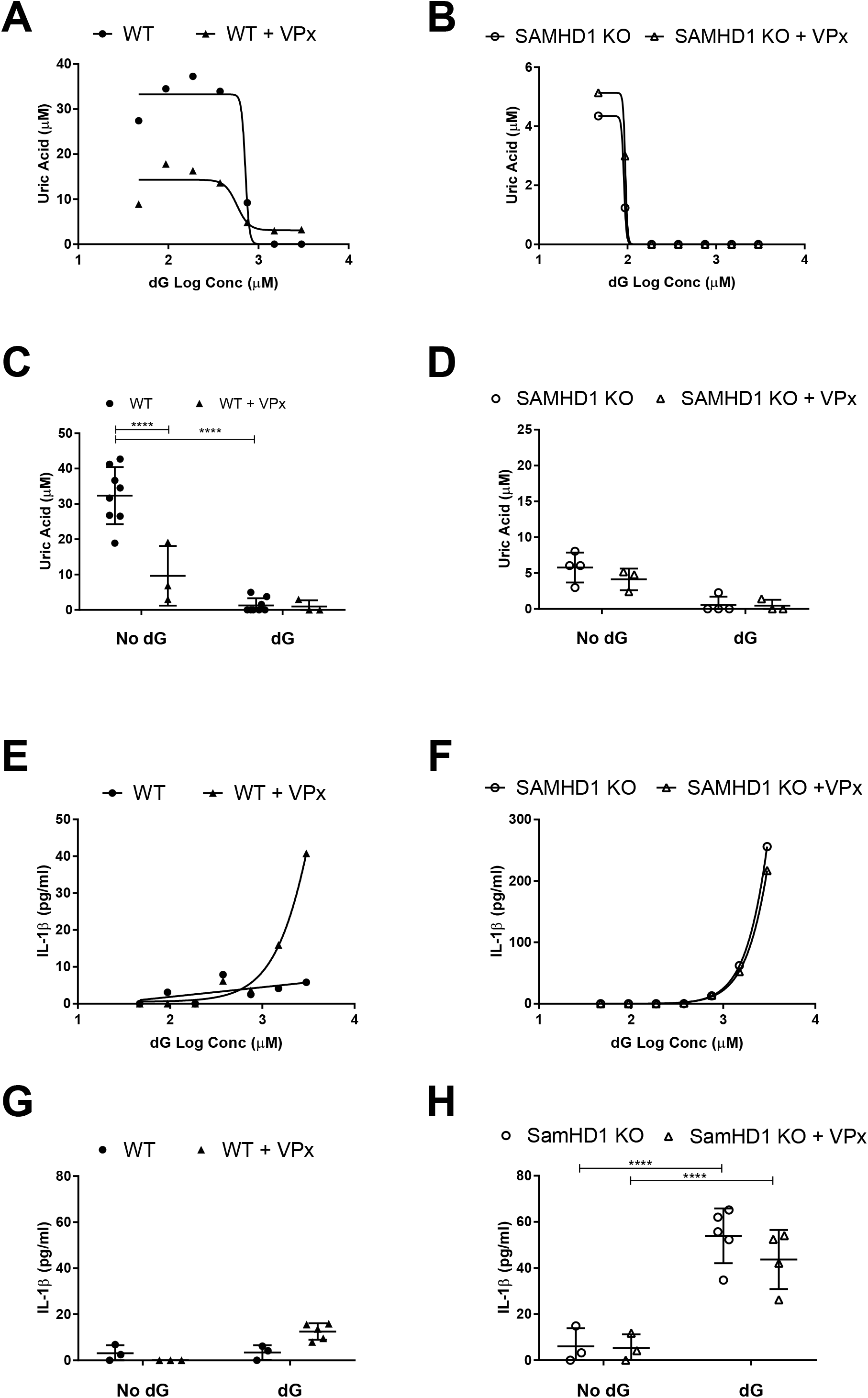
SAMDH1 alters uric acid production and inflammation from macrophages. (A) Uric acid production from wild-type pMac with increasing concentration of dG in the absence (circles) or presence of VLPs carrying VPx (triangles). (B) As in (A) but with SAMHD1 knockout pMac. (C) Uric acid production from wild-type pMac from 3-8 measurements over 3 experiments exposed to no dG or 1.5 mM dG for 4 days in the absence (circle) or presence of VPx (triangles). (****adjusted p<0.0001, 2-way ANOVA with Sidak’s multiple comparison test). (D) As for (C) but with SAMHD1 knockout pMac. (E-H) As in (A-D) but measuring IL-1B release into the supernatant.

### SAMHD1 activity alters the impact of deoxyguanosine on inflammation

Uric acid and crystals of MSU are able to activate macrophages, contributing to the inflammation associated with gout(30). Crystals of MSU are known to activate the NLRP3 inflammasome, thereby activating Caspase 1, resulting in the cleavage of proIL-1β and release of pro-inflammatory cytokine IL-1β(31). As MSU crystals had been observed within wild-type macrophages treated with dG, and we had characterised an increased sensitivity of SAMHD1-knockout pMacs to dG cytotoxicity, we hypothesised that this death would be pro-inflammatory. To detect inflammation, we used an ELISA specific for cleaved IL-1β and observed an increase in IL-1β upon exposure of the pMacs to high levels of dG, but only when SAMHD1 was also inactivated, either with VPx (Figure 4E+G, and Supplementary Figure 2C) or through gene editing (Figure 4F+H). The molecular mechanism of this inflammatory cell death response to elevated dG was investigated by treating SAMHD1 KO pMacs with a low, but toxic, dose of dG in the presence of a variety of inhibitors. Cell death was only affected by two drugs, Necrostatin1 and GSK872, that specifically target RIPK1 and RIPK3, components of the necroptosis pathway (black bars in Supplementary Figure 4)(32), suggesting that dG causes death by necroptosis. However, IL-1β release from the dG-treated cells was only affected by drugs involved in the NLRP3 pathway, i.e the Caspase 1 inhibitors Z-YAD-fmk and Ac-YVAD-CMK, the NLRP3-specific inhibitor, MCC950(33), the NF-κB inhibitor Bay11-7082(34), and the oxidised mitochondrial DNA analogue 8-OH-dG(35) (Supplementary Figure 4 and 5C). The latter suggests a mitochondrial origin of the dG-induced damage that results in IL-1β release, a feature observed in vivo with SAMHD1 deletions(6).

## Discussion

It has become apparent that the deoxynucleotide triphosphohydrolase activity of SAMHD1 affects the efficacy of many nucleoside-based chemotherapeutic agents(12) and mutations in SAMHD1 are linked to the development of cancer(9,10). These two features make the study of SAMHD1 integral to understanding drug susceptibility and resistance of different cancers. In this report we confirm the impact of SAMHD1 on three chemotherapeutic agents using gene edited pluripotent stem cells and expand upon the results by characterizing the effect of SAMHD1 on non-dividing cells. By including the natural nucleoside dG in the panel of drugs, we were able to demonstrate a key link between dNTP metabolism, cell death and inflammation that is not dependent on cell division, but is controlled by SAMHD1. Thus, we have confirmed that SAMHD1 function should be considered as a component of any patient stratification for drug treatment options. We also demonstrate that SAMHD1 is a viable drug target that would act synergistically with nucleoside-based agents in dividing cells and with dG in non-dividing cells. Additionally, our data suggest an alternative hypothesis to the cell type specific loss in ADA and PNP immunodeficiencies. Rather than rapid cell division, per se, being a cause for T cell loss, we would suggest that lack of SAMHD1 buffering of dNTPs within these cells makes them particularly sensitive to imbalances in dNTPs caused by loss of ADA or PNP. If any residual activity of SAMHD1 within these cells could be enhanced (e.g. by inducing its dephosphorylation), then the severity of the disorder may be reduced.

Within dividing cells, we have shown variable effects of SAMHD1 on drug efficacy, depending on stereochemistry at the 2’ position. The importance of this position on the ring is not surprising given that SAMHD1 is inactive against NTPs that contain a 2’OH group. However, the results suggest ways of tailoring future drugs to either be catabolised by SAMHD1 (for enhanced safety when targeting SAMHD1 mutated cancers) or be resistant to SAMHD1 cleavage (for enhanced breadth of activity). The difference in sensitivity to SAMHD1 between Gemcitabine and Ara-C is no doubt key to the narrow suitability of Ara-C compared with the broad range of cancers that Gemcitabine has been successful at treating.

In contrast to their potent activity in diving stem cells, none of the chemotherapeutic agents affected the viability of the non-dividing pMacs, even in the absence of SAMHD1. This would suggest that the processes requiring dNTPs within macrophages, namely DNA repair and mitochondrial DNA replication, are not acutely affected by DNA strand terminators. Conversely the natural nucleoside, dG, did show marked cytotoxicity towards pMacs, but only in the absence of SAMHD1. Although dG has long been known to be toxic, particularly to T cells (e.g. the preparation of alymphoid thymus explants uses dG toxicity to remove thymocytes(36), it had not previously been associated with toxicity in non-dividing cells such as macrophages. Cancer treatment generally focuses on the uncontrollably dividing cancer cells. However, not all cells required for the development of malignant tumours are dividing(26,27). Both quiescent cancer stem cells and TAMs are relatively resistant to standard treatment regimens. Thus, the combination of dG and VPx (or an anti-SAMHD1 drug) would enable the disruption of the tissue microenvironment, which is essential to the survival of the tumour. An additional benefit to this strategy is that we have shown that the cell death that is promoted by dG in SAMHD1-depleted cells is highly inflammatory, which would enhance immunological clearance of the target tumour. Future work should focus on targeted manipulation of SAMHD1 function and alternatives to dG that have improved pharmokinetics but maintain its toxicity and inflammatory consequences.

Deoxyguanosine toxicity was observed exclusively within cells lacking SAMHD1 (e.g. CEM cells, SAMHD1 KO stem cells and VPx-treated macrophages), however this study has provided insights into the homeostatic regulation of purine nucleotides in the presence of SAMHD1. Instead of cell death and inflammation, wild-type cells are able to buffer the effect of exogenous dG, and consequently increased dGTP, through activation of SAMHD1 and allosteric inhibition of RNR by dGTP. Thus, the cell is protected from elevated dGTP, and the intracellular reactions are biased towards catabolism of dGTP, resulting in elevated levels of uric acid. In humans, uric acid is present at very high levels due to the loss of uricase enzyme, which in non-primates converts it into the more soluble allantoin. Uric acid levels within the serum (90-400 μM) are maintained at close to saturation (400 μM) by a combination of production, kidney and gut excretion, and kidney reabsorption. The high levels of uric acid are thought to have beneficial effects due to its antioxidant properties, which have been linked to protection from neurological conditions such as Alzheimer’s disease and Parkinson’s disease(37). Our results show that pMacs excrete a large amount of uric acid at steady state(38). However, we have also shown that this secretion is reduced by inactivating SAMHD1 and converted to storage and crystallisation upon elevation of exogenous dG. How this uric acid secretion occurs and how it would be shut-off are open questions. Within macrophages two of the main uric acid transporters expressed are GLUT9 (SLC2A9) and MRP4 (ABCC4)(39), both of which are detected in the cytoplasm(40,41). We hypothesise that in macrophages, uric acid is routinely transported into vesicles for export. However, by overloading the system with exogenous dG, the vesicles become enlarged (as seen in the TEM images), and upon reaching levels above saturation intra-vesicular crystals form (as see in SEM and TEM images).

The retention of uric acid and formation of crystals upon exposure to exogenous dG could have significant impact within patients suffering from SAMHD1-linked AGS. Within the bone marrow, macrophages phagocytose expelled nuclei from erythroblasts as they terminally differentiate into reticulocytes and during brain development and homeostasis microglia clear apoptotic neurones(42,43). Both these processes would result in high levels of DNaseII-mediated DNA cleavage in the phagolysosmes, releasing nucleosides that in the absence of SAMHD1’s homeostatic regulation, could alter the balance of dNTP metabolism. As shown here in vitro, this imbalance could result in pro-inflammatory cytotoxicity and MSU crystal formation. The localised production of intracellular macrophage crystals would represent a novel damage-associated molecular pattern in the aetiology of AGS. Intriguingly, there is evidence for elevation of IL-1β within the CSF of AGS patients(44), and sufferers often present with the comorbidity lupus(17), an autoimmune disease that also has links to IL-1β(45).

In summary, our results substantiate the notion that SAMHD1 is a pivotal regulator of purine nucleoside metabolism in terminally differentiated cells, and indicates that it might offer a valuable target for therapeutic intervention in a range of inflammatory, infectious and malignant diseases.

### Experimental Procedure

#### Cells

CEM and THP1 cells were cultured in RPMI with 10% FCS and 1% P/S. Pluripotent stem cell lines (Hues2 (WT) and its CRISPR-Cas9 gene edited SAMHD1 KO isogenic clones E2, G10 and F8)(23), OX1.19 (28), SFC841-03-01(46) and SFC856-03-04(47) were all grown on Geltrex (Life Technologies) in E8 (Life Technologies) and passaged with TrypLE (Thermofisher) using Rho Kinase inhibitor Y-27632 (10 μM, Abcam). Stem cells were differentiated into pMac using an established protocol(28). IPS cell lines were originally derived from a healthy donors recruited through the Oxford Parkinson’s Disease Centre having given signed informed consent (Ethics Committee that specifically approved this part of the study, National Health Service, Health Research Authority, NRES Committee South Central, Berkshire, UK, REC 10/H0505/71). All experiments were performed in accordance with UK guidelines and regulations and as set out in the REC.

#### Western Blot

Cells lysates (20 μg per lane) prepared with RIPA buffer (25mM Tris-HCl pH 7.6, 150mM NaCl, 1% NP-40, 1% sodium deoxycholate, 0.1% SDS), containing protease inhibitor cocktail (Sigma) and phosphatase inhibitors (sodium othovanadate, sodium fluoride and beta-glycerophosphate) were separated on NuPAGE Bis-Tris gels (Thermofisher) and transferred onto PVDF membranes. Membranes were probed with mouse anti-SAMHD1 (2D7, Origene, 1:2000), rabbit anti-phT592-SAMHD1 (Cell Signalling Technology, 1:1000) and rabbit anti-Tubulin (Sigma,1:5000).

#### Cell viability assay

Cells were plated in 96 well plates at 3×10^5^ (THP1, CEM and stem cells) or 5×10^4^ cells per well (pMacs) in 100 μl media. The following day drug titrations were added with or without pre-incubation for 2 hours with 7.5 ng per well SIV-VLP containing VPx. Forty-eight hours (96 hours for experiments with pMacs) post addition of drugs the cells were assayed for viability using the CellTiter-Fluor™ (Promega) following the manufacturers protocol.

#### Uric Acid and IL-1β assays

Supernatants from 5×10^4^ macrophages in 96 well plates were harvested and assayed for uric acid levels using a fluorescent based assay (Cayman chemicals) following the manufacturers protocol. Where necessary uric acid was extracted from the cells by incubation with 1M KOH for 5 minutes followed by neutralisation with 1M acetic acid. The same supernatants were also assayed for IL-1β levels by ELISA (ThermoFisher Scientific).

#### TEM protocol

Cells plated on Thermonox coverslips were fixed with pre-warmed fixative (2.5% glutaraldehyde, 2% PFA in 0.1M PIPES buffer, pH 7.2). Cells were washed in 0.1M PIPES buffer pH 7.2 for 3x 15 mins, then in 50 mM glycine in 0.1M PIPES buffer, pH 7.2 for 20 mins. Cells were then incubated in 1% osmium tetroxide in 0.1M PIPES buffer, pH 7.2 for 1 hr at 4°C, then washed with water for 60 mins and incubated overnight in 0.5% uranyl acetate (aqueous) at 4 °C. Following a water wash, samples were dehydrated (30%, 50%, 70%, 80%, 90% and 95% ethanol for 10 mins each, then 100% ethanol for 3x 30 mins) and then gradually infiltrated with Agar100 epoxy resin, starting with 25% resin for 1 hrs, 50% resin for 2 hrs, 75% resin for 1 hr and 100% resin overnight, followed by two changes with fresh 100% resin the next day before embedding by inverting the coverslip (cell side down) onto a tube filled with fresh resin. Blocks were polymerised for 48 hrs at 60 °C, then submerged in liquid nitrogen and the coverslip snapped off, to leave the cells embedded as a monolayer in the resin. Ultrathin (90 nm) sections were obtained using a Leica UC7 ultramicrotome with a diamond knife (Diatome) and placed on 200 mesh or formvar-coated 50 mesh Copper grids, then post-stained with Reynold’s lead citrate for 5 min, and imaged on a FEI Tecnai 12 TEM operated at 120 kV using a Gatan OneView CMOS camera.

#### SEM protocol

As for the TEM protocol up to the end of the dehydration series, except that the glycine wash and UA staining steps were omitted. After the final 100% ethanol incubation, cells were treated with 100% hexamethyldisilazane for 3mins and then air dried. Samples were sputter coated with a ∼10nm gold layer and imaged with a Zeiss Sigma 300 FEG-SEM at an accelerating voltage of 2kV.

#### Birefringence

Cells were fixed as for EM analysis, then dehydrated twice for 20 seconds in 100% ethanol before being stained for 20 seconds in alcoholic Eosin Y (Leica Biosystems). Cells were then destained with two 20 second washes in 100% ethanol and two 20 second washes in xylopath before DPX mounting. Images were taken with a Nikon Optiphot-Pol Petrological microscope with transmitted and incident illumination using a gypsum plate red filter.

## Figure Legends

**Sup Fig 1.**
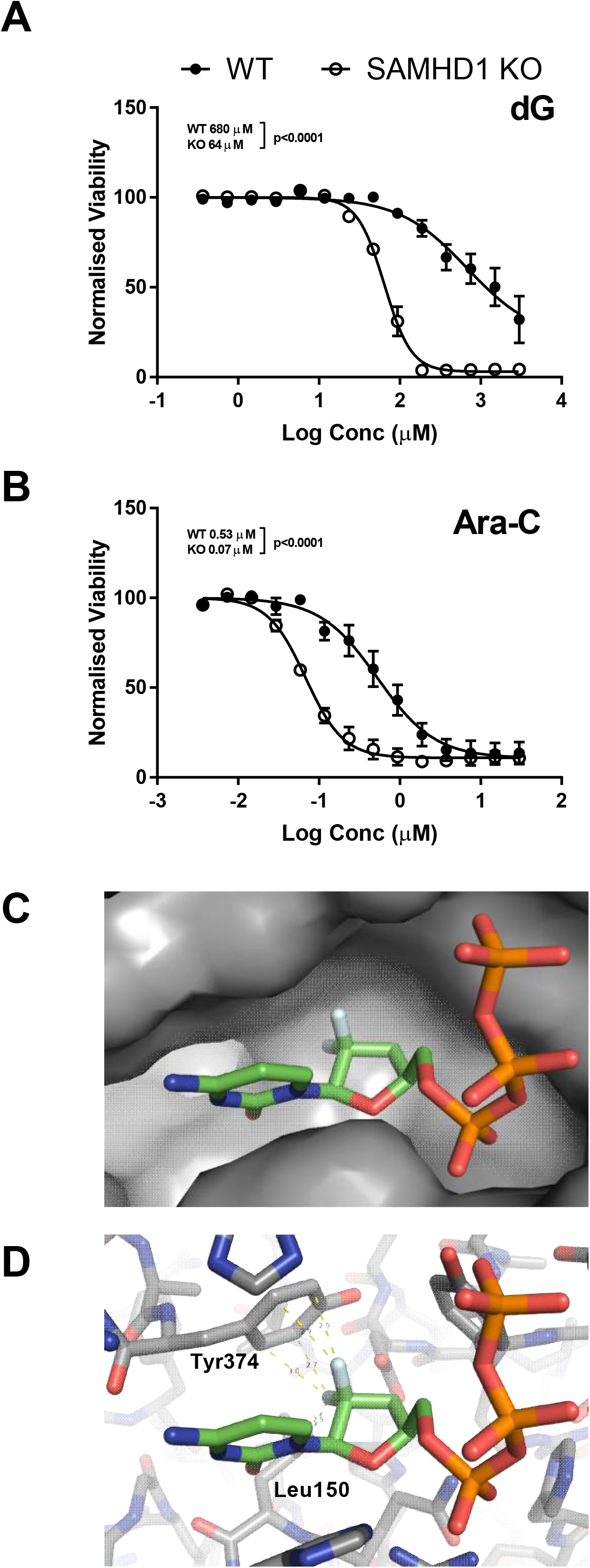
Effect of SAMHD1 on dG and Ara-C toxicity to stem cells is not dependent on genotype. Three individual donor-derived pluripotent stem cell lines (Hues2, OX1.61 and 856; WT) and three separate clones of SAMHD1 knockout lines derived from the Hues2 WT line (E2, G10 and F8; SAMHD1 KO) were compared for their sensitivity to (A) dG and (B) Ara-C. Results are plotted as mean +/- SD of the relative viability to an untreated control sample. P-values represent the statistical significance for the EC50 measurements being different (F-test). (C) Model of a di-fluorinated nucleotide (i.e. gemcitabine) within a surface render of the active site of SAMHD1. The structure is based on PDB-ID 4to4, with the drug rendered as sticks. (D) Stick representation of both di-fluorinated nucleotide and SAMHD1 showing the distances between the fluorines (in silver) and Leu150 (2.5 Ångströms) and Tyr374 (2.7-3.0 Ångströms).

**Sup Fig 2.**
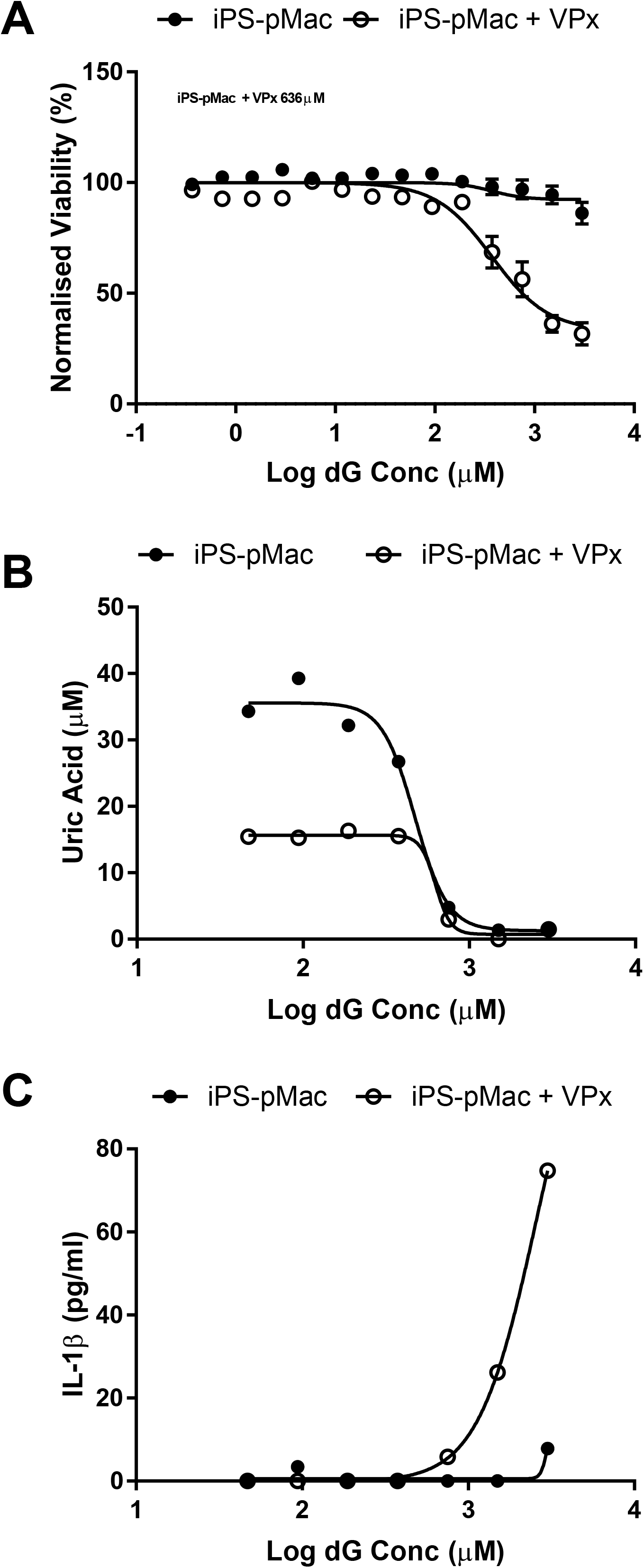
Deoxyguanosine toxicity is SAMHD1 dependent and results in loss of uric acid production but release of IL-1B. (A) Viability assays for pMacs differentiated from wild-type induced pluripotent stem (iPS) cell line OX1.19 (closed circles) or exposed to VLPs carrying VPx (open circles) in the presence of increasing concentrations of dG. Results are plotted as mean +/- SD of the relative viability to an untreated control sample. (B) As for (A) but assaying the supernatant for uric acid release in a single titration. (C) As for (B) but assaying for IL-1β release.

**Sup Fig 3.**
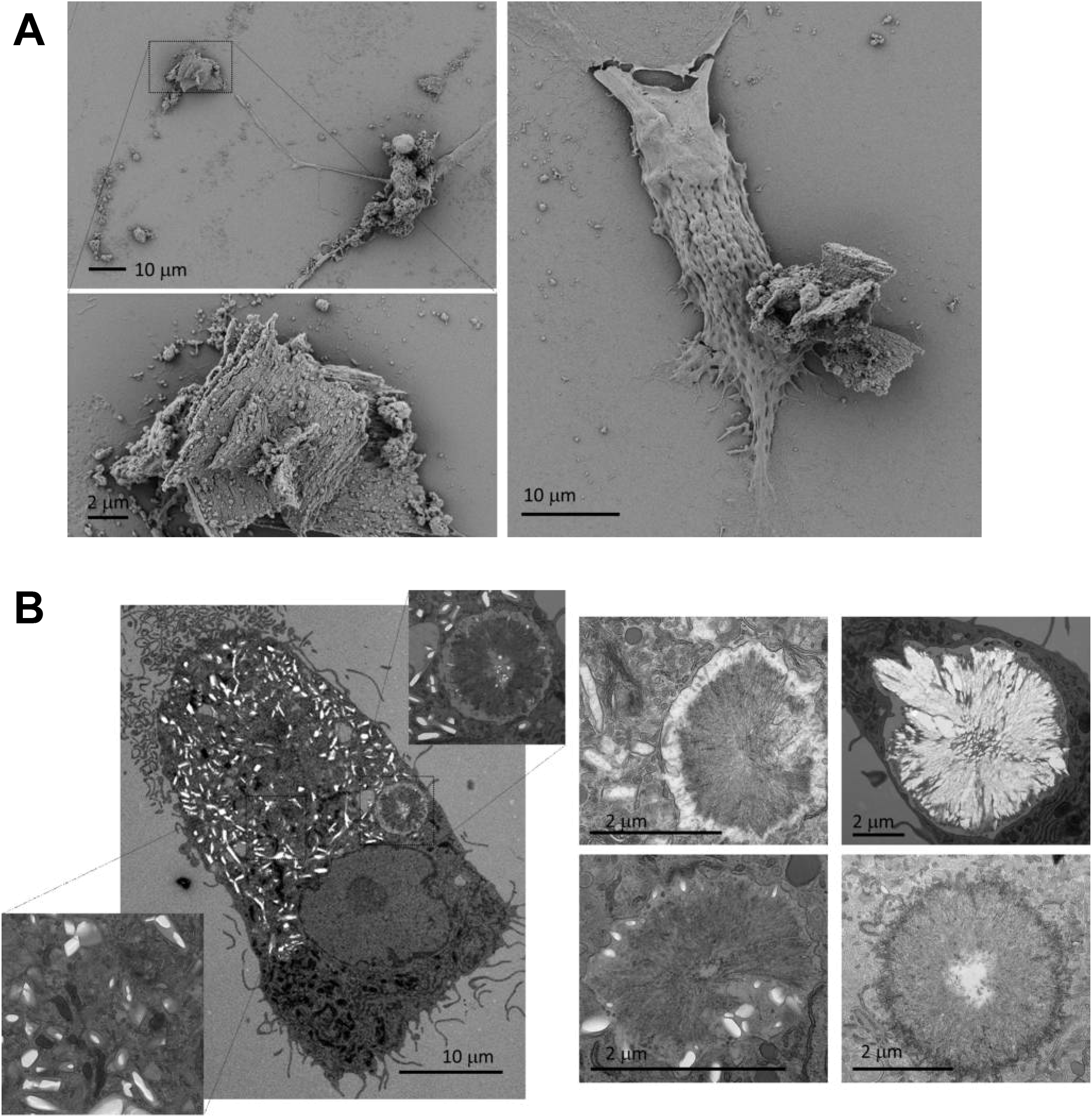
Deoxyguanosine treatment of wild-type macrophages results in crystal formation. (A) SEM of wild-type pMac exposed to 1.5 mM dG for 8 days. The bottom left figure is highlighting the crystalline plates within the cellular debris. (B) TEM images of monocyte-derived macrophages treated with 1 mM dG for 7 days showing multiple morphological alterations caused by dG treatment.

**Sup Fig 4.**
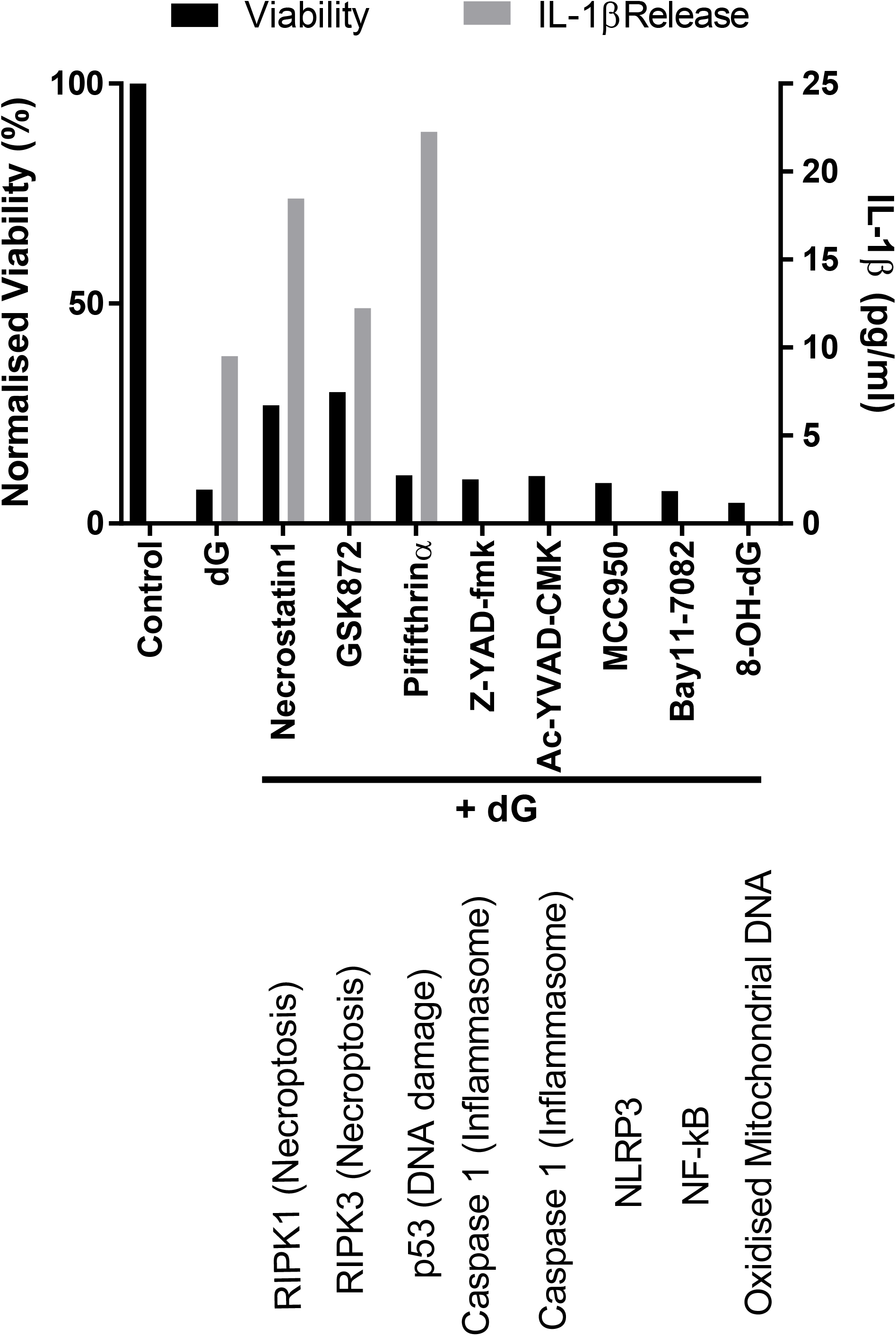
Deoxyguanosine causes necroptosis and NLRP3 inflammasome activation. Cell viability (black bars, left y-axis) and IL-1β release (grey bars, right y-axis) from SAMHD1 knockout pMac cells treated with 0.3 mM dG with the indicated inhibitors for 4 days.

**Sup Fig 5.**
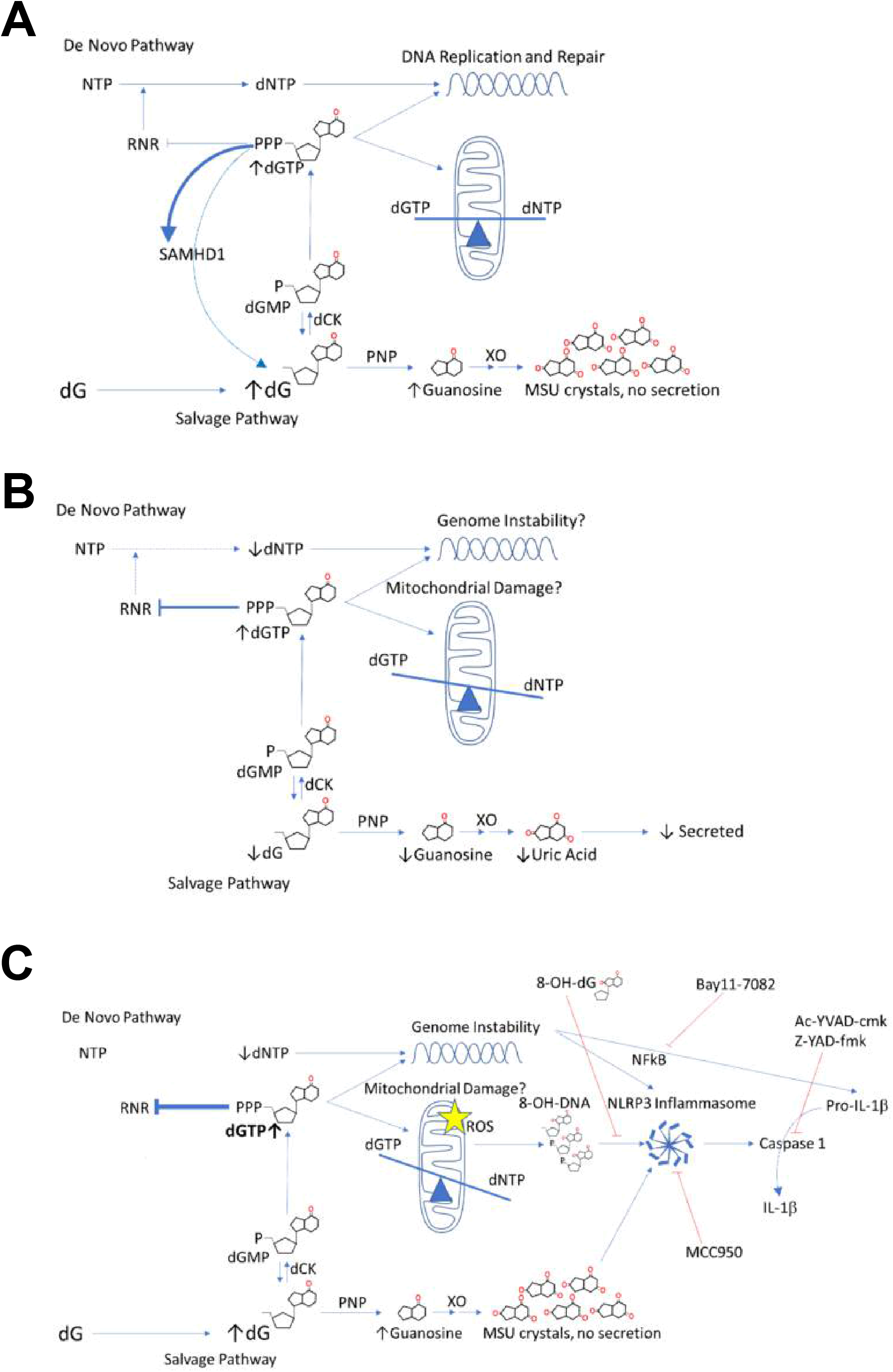
Models of dGTP metabolism in (A) wild type cells under conditions of excess dG, (B) SAMHD1 KO cells under physiological conditions and (C) SAMHD1 KO cells under conditions of excess dG. Abbreviations: rNTP ribonucleotide triphosphate, dNTP deoxynucleotide triphosphate, dG deoxyguanosine, RNR ribonucleotide reductase, dCK deoxycytidine kinase, PNP purine nucleoside phosphorylase, XO xanthine oxidoreductase, MSU monosodium urate.

## Acknowledgments

This publication arises from research funded by the John Fell Oxford University Press (OUP) Research Fund. The iPS cell line used in this study was originally generated from a donor sample supplied by the Oxford Parkinson’s Disease Centre (OPDC) study (funded by the Monument Trust Discovery Award from Parkinson’s UK, a charity registered in England and Wales (2581970) and in Scotland (SC037554), with the support of the National Institute for Health Research (NIHR) Oxford Biomedical Research Centre based at Oxford University Hospitals NHS Trust and University of Oxford, and the NIHR Comprehensive Local Research Network), and was reprogrammed within StemBANCC, (supported by the Innovative Medicines Initiative Joint Undertaking under grant agreement number 115439, resources of which are composed of financial contribution from the European Union’s Seventh Framework Program (FP7/2007e2013) and EFPIA companies’ in kind contribution).

## Author Contributions

MDM designed, carried out, analysed data and prepared the manuscript. EJ and CJ carried out experiments. OG generated polarised microscopy images. SJ performed in silico modelling. WJ analysed data and prepared the manuscript.

